# Multilevel selection in the evolution of sexual dimorphism in phenotypic plasticity

**DOI:** 10.1101/2022.12.16.520722

**Authors:** Anton S. Zadorin, Olivier Rivoire

## Abstract

Phenotypes are partly shaped by the environment, which can impact both short-term adaptation and long-term evolution. In dioecious species, the two sexes may exhibit different degrees of phenotypic plasticity and theoretical models indicate that such differences may confer an adaptive advantage when the population is subject to directional selection, either because of a systematically varying environment or of a load of deleterious mutations. The effect stems from the fundamental asymmetry between the two sexes: female fertility is more limited than male fertility. Whether this asymmetry is sufficient for sexual dimorphism in phenotypic plasticity to evolve is, however, not obvious. Here, we show that even in conditions where it provides an adaptive advantage, dimorphic phenotypic plasticity may be evolutionarily unstable due to sexual selection. This is the case, in particular, for panmictic populations where mating partners are formed at random. However, we show that the effects of sexual selection can be counteracted when mating occurs within groups of related individuals. Under this condition, sexual dimorphism in phenotypic plasticity can not only evolve but offset the twofold cost of males. We demonstrate these points with a simple mathematical model through a combination of analytical and numerical results.

## I. INTRODUCTION

Phenotypic plasticity, the influence of the environment on the development of phenotypes, is a well-recognized component of evolutionary change [1] with the potential to constitute an essential adaptation to varying environments [2]. Sex differences in phenotypic plasticity are documented in several dioecious species [3, 4]. Although considerable variation is observed between species, the general trend is that females are more plastic than males. This trend is, for example, observed in an analysis of body mass plasticity in insects where, in addition, the average plasticity appears to be greater in females [3]. Similarly, a meta-analysis of data on sex-specific plasticity in response to thermal adaptation revealed that of seven categories of traits, one showed a significant difference in plasticity between the sexes, namely cold resistance, which appears to be more plastic in females [4]. Only recently, however, has the question of the adaptive value of these sex differences started to be studied mathematically [4, 5]. Remarkably, these studies show that sexual dimorphism in phenotypic plasticity can enhance population growth and thus prevent the extinction of populations subject to directional selective pressures. They have left open, however, the question of the conditions under which such an adaptive dimorphism may evolve [4, 5]. A puzzling observation is indeed that sexual dimorphism in phenotypic plasticity can fail to evolve even in conditions where it is adaptive [5].

Understanding the conditions under which sexual dimorphism in phenotypic plasticity can evolve has at least two motivations. On one hand is the pressing issue of determining the role that phenotypic plasticity may play in the adaptation of extant species to current climate changes [6]. On the other hand is the long-standing question of the origin and prevalence of dioecious species given the “two-fold cost of males” that they incur compared to monoecious species: a monoecious species where each individual would be as fecund as the females of a dioecious species would indeed produce twice as many offsprings per generation and therefore have a major selective advantage [7]. This raises two questions: (1) Under which conditions may sexual dimorphism in phenotypic plasticity be an adaptation that offsets the two-fold cost of males? (2) Under which conditions may such a population-level adaptation be reachable through evolution?

In previous work [5], we started to examine these questions with a focus on the adaptive value and evolution of sexual dimorphism in developmental variances – the possibility that one sex displays higher developmental canalization than the other. We found that such a sexual dimorphism in phenotypic plasticity could in principle be adaptive but only under restricted conditions that made it unlikely to be a generic cause of phenotypic differences between sexes, or a fortiori to explain why many species comprise two sexes. Further, we found that an optimal difference in developmental variances failed to evolve even in conditions where it was adaptive. Here we revisit these questions with a focus on sexual differences in phenotypic plasticity. Although the two problems can formally be mapped on each other [5], we show that sexual dimorphism in phenotypic plasticity can possibly confer a substantial adaptive advantage over a much broader range of conditions.

To demonstrate this result, we study a simple but generic model for the evolution of sexually dimorphic plasticity. This model extends previous models [2, 4, 5, 8, 9] with several crucial differences. First, it differs from models of quantitative genetics [2, 4, 8] by introducing an explicit dynamic for genetic changes, which allows us to readily analyze the evolution of plasticity through modifier genes [10]. Second, compared to our previous models [5, 9], it accounts for varying degrees of sex-specific limitations in fertility, as well as for the possibility that mating interactions are limited to groups of genetically related individuals. We show that mating patterns – the rules that govern the associations between males and females – are critical to both the adaptive value and the evolution of dimorphic plasticity. In particular, we identify simple mating patterns for which sexual dimorphism in phenotypic plasticity is evolvable with an adaptive value that offsets the two-fold cost of males.

## II. MODEL

### A. Model with non-evolving plasticity

We consider a population of individuals, each with a continuous one-dimensional genotype *γ*, a continuous one-dimensional phenotype *ϕ* and a sex •, which can either be female (• = **♀**) or male (• = **♂**). At each discrete generation *t*, the environment *x*_*t*_ defines an optimal phenotype, which may be directionally varying as *x*_*t*_ = *c*_*E*_*t*, where *c*_*E*_ therefore represents a rate of change. The phenotype *ϕ* of an individual dictates its chances of survival and reproduction. Following previous models [2, 4, 5, 8, 9], we assume that the probability for an individual with phenotype *ϕ* to survive and possibly reproduce in environment *x* is 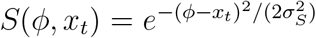 where 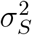 represents the stringency of selection. We assume that an individual whose parents have genotypes *γ*^♀^ and *γ*^♂^ inherits a mid-genotype

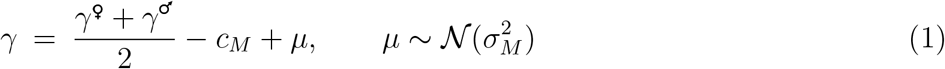

where *µ* is a random variable normally distributed with zero mean and variance 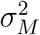 and where *c*_*M*_ is a possible mutational bias towards deleterious mutations when considering *c*_*M*_ > 0. This form of inheritance is justified if *γ* represents the contribution of many loci that are independently transmitted at random from each parent.

Following previous models [2, 4, 5, 9], we consider the phenotype *ϕ* of an individual of sex • to possibly depend both on its genotype *γ* and on its environment *x*_*t*_ through the relation

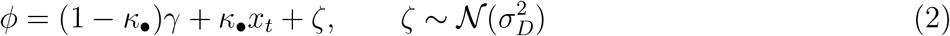

where 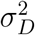 represents a developmental variance. *κ*_•_ is interpreted as a reaction norm, which may here depend on the sex • of the individual. We previously examined the case of sex-specific developmental variances [5] but assume here that a common 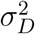 applies to both sexes.

Finally, the sex of an individual is chosen at random with no bias, • = **♀** or **♂** with same probability, which we write as • = rand([**♀, ♂**]).

### B. Model with evolving plasticity

Instead of treating the reactions norms *κ*_♀_ and *κ*_♂_ as common non-evolving parameters that are set to given values for all members of the population, we may treat them as evolving traits controlled by genotypic variables, which may differ between members of the population. To this end, the reaction norms can either be integrated in the genotypes or viewed as phenotypes that are controlled by new variables introduced in the genotypes. We adopt this second formulation so that all components in the genotype are continuous variables subject to Gaussian mutational effects. We therefore introduce in the genotype modifier genes *y*_♀_, *y*_♂_ that respectively control the female and male reaction norms *κ*_♀_ and *κ*_♂_. The complete genotype of an individual is of the form Γ = (*γ, y*_♀_, *y*_♂_); these variables are a priori independent but typically become coupled through evolution. In this more general model, the phenotype of an individual consists of its continuous trait *ϕ*, its sex • and its reaction norm *κ*_•_. It has therefore the form Φ = (*ϕ*, •, *κ*_•_). *κ*_•_ is directly related to *y*_•_ by

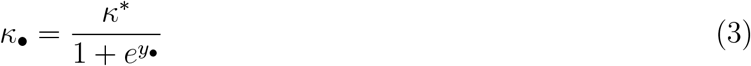

so as to map the continuous trait *y*_•_ into an interval [0, *κ*^*^] where *κ*^*^ < 1 represents a maximal reaction norm.

We assume that the modifier genes follow the same inheritance rules as *γ*, i.e., given parents with genotypes 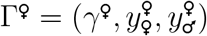 and 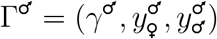, we assume that an offspring inherits

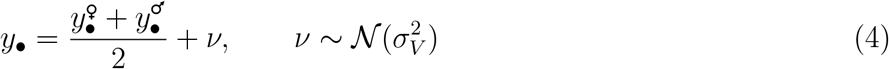

with • = **♀** and **♂**. Here we introduce a mutational variance 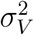 that may differ from 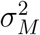 but note that we could equivalently consider 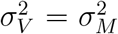 by redefining *κ*_•_ as 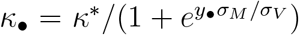. The present formulation is adopted to recover the case of a non-evolving plasticity as the limit where 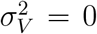, in which case *y*_•_, and therefore *κ*_•_, does not change. For simplicity, we also assume here no mutational bias.

The introduction of a maximal reaction norm *κ*^*^ < 1 is motivated by the fact that in absence of further constraints, *κ*_•_ = 1 guarantees that *ϕ* = *x*_*t*_, i.e., perfect adaptation independently of the genotype *γ*. This extreme degree of plasticity is optimal but not biologically feasible and models usually integrate a fitness cost that penalizes large values of *κ*_•_ [2]. Our parameter *κ*^*^ is a simple representation of this cost, with values *κ* ≤ *κ*^*^ corresponding to no cost and values *κ* > *κ*^*^ to infinite cost.

### C. Mating patterns

Following Eqs. (2)-(3), a newborn individual with genotype Γ = (*γ, y*_♀_, *y*_♂_) develops to have phenotype Φ = (*ϕ*, •, *κ*). This individual reaches maturation with probability 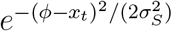 and otherwise dies. Next, we assume that each mature individual of sex • = **♀** or **♂** has the capacity to participate in *q*_•_ mating events, each producing one individual of the next generation. This maximal capacity may, however, not be achieved if partners of the opposite sex are not available. If *M*_♀,*t*_ and *M*_♂,*t*_ are, respectively, the number of mature females and males at generation *t*, the number of newborn at generation *t* + 1 is therefore

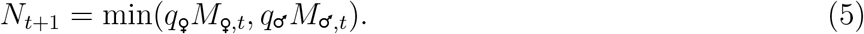

This relationship accounts both for the case where the limitation is the number of females (*q*_♀_*M*_♀,*t*_ < *q*_♂_*M*_♂,*t*_) or the number of males (*q*_♀_*M*_♀,*t*_ *> q*_♂_*M*_♂,*t*_). We consider females as the limiting sex, *q*_♀_ < *q*_♂_, with a limit case being males having unlimited mating capacities (*q*_♂_ = ∞) so that *N*_*t*+1_ = *q*_♀_*M*_♀,*t*_.

We also consider the possibility for mating to occurs only within groups. We take these mating groups to have a maximal size *K* and assume that when the number of newborn individuals within a group exceeds *K*, this group splits into randomly formed sub-groups of same size. This structure into groups is only relevant to the choice of mates while viability and density selection (the resampling of the population to reach the imposed total size *N*) are performed at the level of the entire population (Fig. 1; see aso Methods for an extension with density regulation at the level of groups and migration between groups). A limit case here is that of panmictic populations where couples are formed totally at random between available mature individuals of each sex, which corresponds to *K* being equal or larger than the population size.

**FIG. 1:**
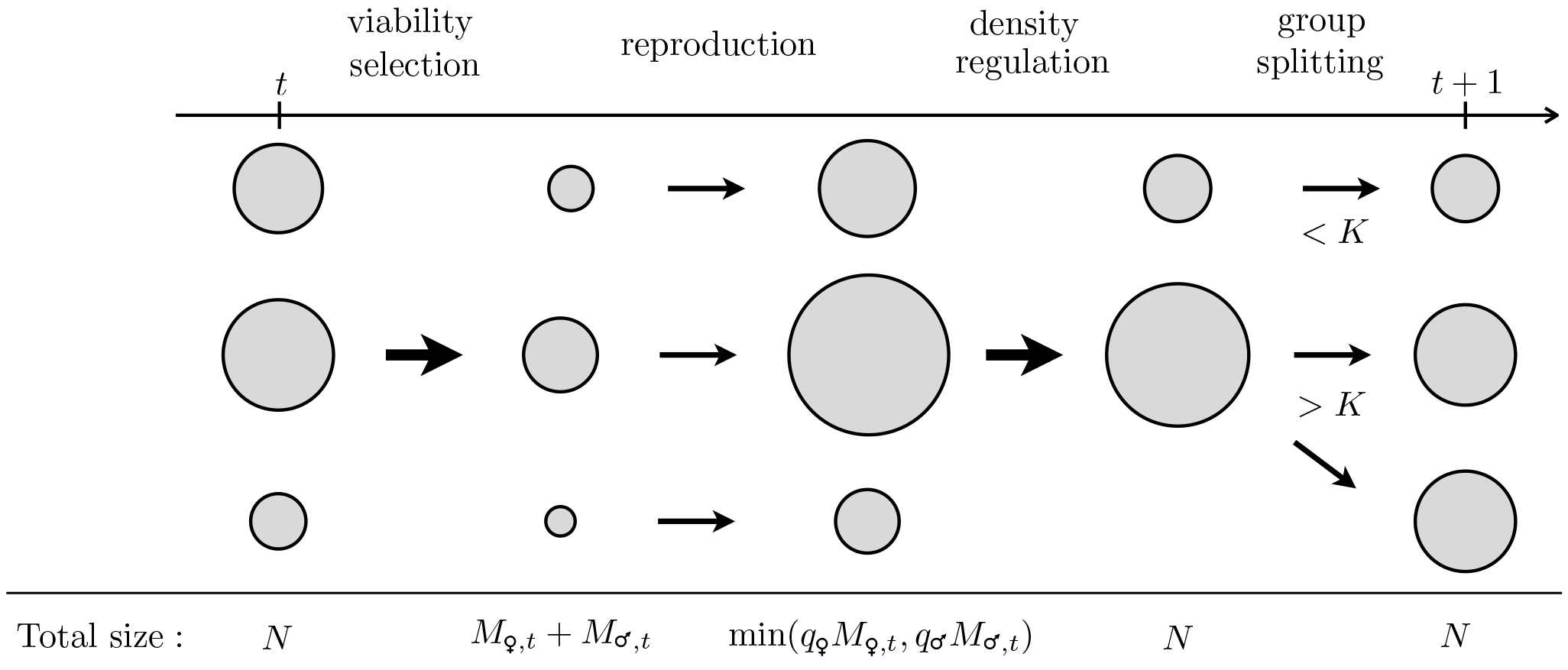
Life cycle in the model – At generation *t*, the population initially consists of *N* newborns divided (or not) into different mating groups. First, depending on its phenotype (including its sex) and on the environment but not on its group, each individual either maturates or dies (viability selection). Second, sexual reproduction occurs between individuals of opposite sex belonging to the same group. Third, the total population size is brought back to *N* by randomly sampling *N* individuals irrespectively of their group but without changing their group identity. Finally, if the size of a group exceeds a given threshold *K*, it is randomly divided into two groups of equal size, thus defining the population of *N* newborns at generation *t* + 1. In particular, if *K* ≥ *N* and the population initially consists of a single group, no group is ever formed and the population is panmictic. The largest arrows indicate the steps that are independent of the structure into groups and the smaller arrows those that are dependent on it. The model can be extended to include group-specific density regulation (soft-selection) or/and migration between groups (Methods).

In general, we expect a population following these rules to grow exponentially whenever *q*_♀_ and *q*_♂_ are sufficiently large, with a growth rate Λ such that *N*_*t*_/*N*_0_ ∼ *e*^Λ*t*^ for large *t*. This growth rate can be computed analytically for large panmictic populations (Methods). More generally, the dynamics can be studied numerically with an agent-based simulation. In practice, it is convenient in numerical simulations to impose a total population size *N* and resample individuals at each generation to maintain this size (Methods). For panmictic populations, we can then compare the analytically computed growth rate Λ to its numerical approximation 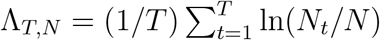 where *N*_*t*_ is the number of newborn individuals at generation *t* and *T* the total number of generations.

### D. Parameters

As so far defined, the model has ten parameters, 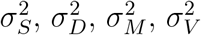, *c*_*E*_, *c*_*M*_, *κ*^*^, *q*_♀_, *q*_♂_, *K* (see Table I), not counting the initial conditions, the total population size *N* and the total number of generations *T* that must also be set when performing numerical simulations. Our results, however, depend only on a subset of these parameters and we therefore set the others. For instance, by rescaling other variances we can always assume 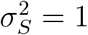 [9]. When *q*_♂_ = ∞, *q*_♀_ contributes only additively to the growth rate and as we are only interested in differences in growth rates, we can also fix *q*_♀_. Additionally, we note that *c*_*E*_ and *c*_*M*_ play formally equivalent roles and can therefore be represented by a single parameter *c* which quantifies a directional selection independently of its environmental or genetic origin (Methods).

**TABLE I:** Summary of the main notations used in the text and figures.

|  |  |
| --- | --- |
| Evolving variables |  |
| $\bullet$ | sex, which may be either female ($\bullet = \text{♀}$) or male ($\bullet = \text{♂}$) |
| $\phi$ | one-dimensional trait under direct selection |
| $\gamma$ | genotype controlling the main trait $\phi$ |
| $\kappa_\bullet$ | sex-specific reaction norm |
| $y_\bullet$ | modifier gene controlling the reaction norm $\kappa_\bullet$ |
| Environmental parameters |  |
| $x_t$ | environmental variable defining the optimal phenotype $\phi$ at generation $t$ |
| $c_E = c$ | change in optimal phenotype per generation ($x_t = c_E t$ after $t$ generations) |
| $\sigma_S^2$ | stringency of selection |
| Genetic parameters |  |
| $\kappa^*$ | maximal reaction norm |
| $c_M = c$ | mean effect of deleterious mutations |
| $\sigma_M^2$ | mutation rate for trait $\gamma$ |
| $\sigma_V^2$ | mutation rate for the modifier genes $y_\text{♀}$ and $y_\text{♂}$ ($\sigma_V^2 = 0$ for non-evolving reaction norms) |
| $\sigma_D^2$ | developmental variance |
| Mating parameters |  |
| $q_\bullet$ | maximal number of matings to which an individual of sex $\bullet$ can participate |
| $K$ | maximal size of the groups within which mating is restricted |
| Population parameters |  |
| $N$ | total population size |
| $\Lambda$ | long-term population growth rate |
| $\chi_\bullet$ | fraction of individuals of sex $\bullet$ reaching maturation at each generation |

We first examine the model with non-evolving reaction norms 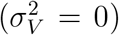 in the limit of an infinite panmictic population (*N* = *K* = ∞). In this case, two additional parameters are the non-evolving reaction norms of each sex, *κ*_♀_ and *κ*_♂_. Next, we consider the long-term evolution of large populations with 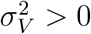, in which case *κ*_♀_ and *κ*_♂_ can differ between individuals of a same population, thus leading us to analyze population-wise mean values 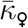 and 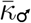.

Unless otherwise indicated, we take the following values of the parameters: *q*_♂_ = ∞, *q*_♀_ = 4, *c* = 10^−1^, 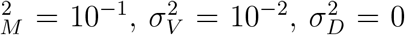, *κ*^*^ = 0.5, and, when performing numerical simulations, *N* = 10^3^ and *T* = 10^4^ with initial conditions such that *κ*_♀_ = *κ*_♂_ = *κ*^*^/2 for all individuals, which uniquely determines the initial conditions for *y*_♀_ and *y*_♂_ by Eq. (3). Finally, we initialize all individuals with *γ* = 0.

## III. RESULTS

### A. Optimal plasticity

For infinite panmictic populations (*N* = *K* = ∞) with non-evolving reaction norms 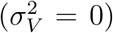, the long-term growth rate Λ can be computed analytically (Methods), with results matching well with those of numerical simulations where *N* = *K* = 10^3^ (Fig. 2A). These results indicate that the growth rate Λ is maximal when the two sex-specific reaction norms *κ*_♀_ and *κ*_♂_ take different values, namely 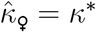 and 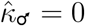 (Fig. 2B). This corresponds to maximally plastic females and minimally plastic males, consistent with previous findings showing that sexual dimorphism in phenotypic plasticity can optimize the longterm growth of populations subject to directional selection [4, 5]. Given that fertility is limited by females, this result may be interpreted as follows: maximizing female plasticity maximizes the number of offspring at the next generation, while minimizing male plasticity maximizes the fitness value of the genotype that these offsprings inherit, as only males with well adapted genotypes reach maturity. In other words, males pay the price of natural selection to guarantee long-term adaptation while females buffer environmental effects to guarantee short-term growth. These results depend primarily on the assumption that fertility is limited by females (*q*_♂_ » *q*_♀_) and the opposite conclusions are reached if *q*_♂_ « *q*_♀_. When *q*_♀_ < *q*_♂_ but *q*_♂_ is comparable to *q*_♀_, we also find that the optimal female reaction norm 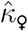 takes a non-zero value that is independent of the maximal possible reaction norm *κ*^*^ (Fig. 2C).

**FIG. 2:**
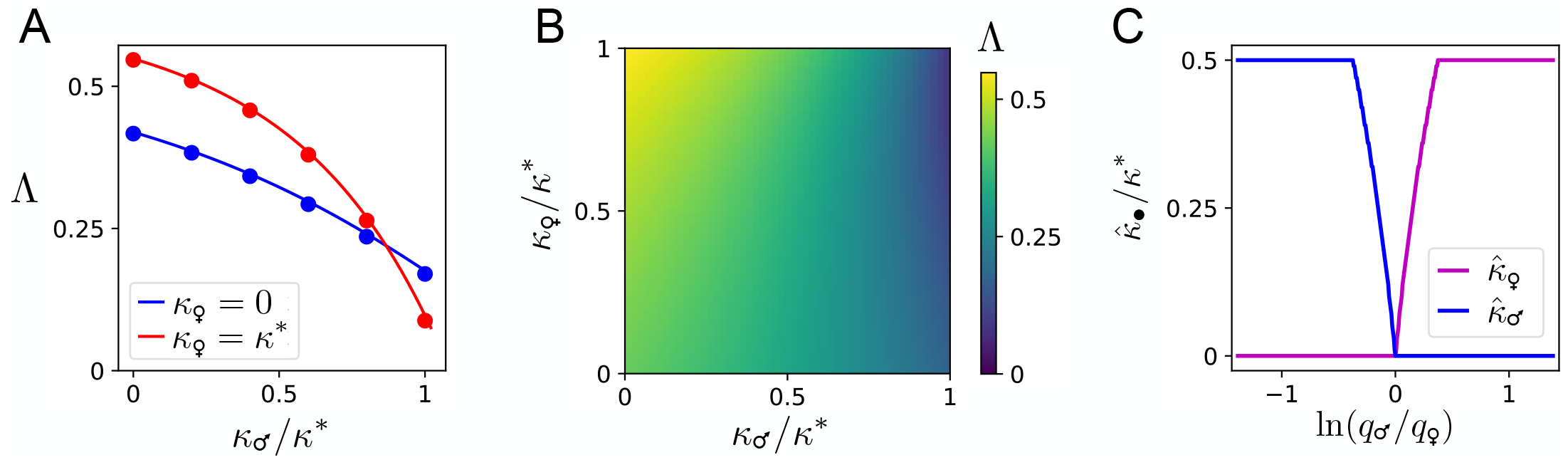
**A**. Long-term growth rate Λ of large unstructured populations with non-evolving reaction norms as a function of the male reaction norm *κ*_♂_ for two extreme values of the female reaction norm *κ*_♀_. The full lines are from analytical calculations and the dots from numerical simulations, showing very good agreement. **B**. Long-term growth rate Λ as a function of *κ*_♂_ and *κ*_♀_, showing a maximum for dimorphic plasticity when *κ*_♂_ = 0 and *κ*_♀_ = *κ*^*^ (upper left corner). **C**. Values of the reaction norms that optimize the growth rate Λ as a function of the ratio *q*_♂_/*q*_♂_ where *q*_•_ controls the maximal number of mating events to which an individual of sex • can participate (• = ♀ or ♂). The case of female demographic dominance corresponds to *q*_♂_/*q*_♀_ > 1, in which case it is optimal for females to be plastic 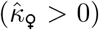 and for males to be non-plastic 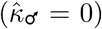. For sufficiently large ratios *q*_♂_/*q*_♀_, it is even optimal for female reaction norms to be maximal (*κ*_♀_ = *κ*^*^). The conclusions are symmetrically reversed if *q*_♂_/*q*_♀_ < 1.

### B. Evolution of plasticity

While a dimorphic plasticity with *κ*_♀_ = *κ*^*^ and *κ*_♂_ = 0 maximizes long-term population growth, we find that when reactions norms are subject to evolution 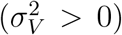, they converge to the same value, namely 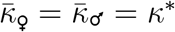 (Fig. 3A). Evolution thus leads to monomorphic plasticity that is sub-optimal for long-term growth. This may be interpreted as arising from a conflict between two levels of selection: long-term population growth benefits from low male plasticity but sexual selection among males promotes high male plasticity. The outcome of such a conflict can be reversed by group selection [11], as we verify here by considering mating to be structured into groups. We find that in presence of mating groups, evolution of the reaction norms can indeed lead to a dimorphic plasticity that maximizes population growth (Fig. 3C). This requires the maximal size of the groups to be sufficiently small (Fig. 3D). This critical size *K*^*^ is found to scale with the total population size *N* as *N*^1/2^ (Fig. S1), which indicates that sexual dimorphism in plasticity can evolve in presence of larger groups when the total population size is itself larger. We also verify that *K*^*^ scales with 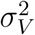 as 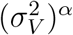 (Fig. S1), as is typical for this type of models [12]. Limiting mating interactions to individuals from a same group enables the evolution of dimorphic plasticity whenever fertility is controlled by females, even if male fertility is also limited, provided this limitation is to a lesser extent (Fig. 3E). When the two sexes have the same fertility limits, however, no asymmetry is present and we verify that no dimorphism evolves (Fig. 3F). The underlying mechanism is common to other problems where a division of the population into groups leads to the evolution of a cooperative behavior that is counter-selected in absence of groups, provided sufficient variance is present between groups [11]. As the mating groups comprise genetically related individuals, this is also the same mechanism as in models based on kin selection [11]. However, one difference with most models involving group selection or kin selection is that both viability selection and density selection are independent of the group in our model: only the choice of a mate depends on it (Fig. 1). The model can be extended to include density regulation at the group level, thus interpolating between two limits known as hard and soft selection [13] (Methods). While a density regulation confined to each group (pure soft selection) prevents any form of group selection [14], we verify that limited density regulation at the group level does not affect our conclusions (Fig. S3A). These conclusions are also robust to the presence of moderate migration between groups (Fig. S3B). Taking these two extensions into account, both group-level density regulation and migration must be sufficiently weak for sexual dimorphism in phenotypic plasticity to evolve (Fig. S3C).

**FIG. 3:**
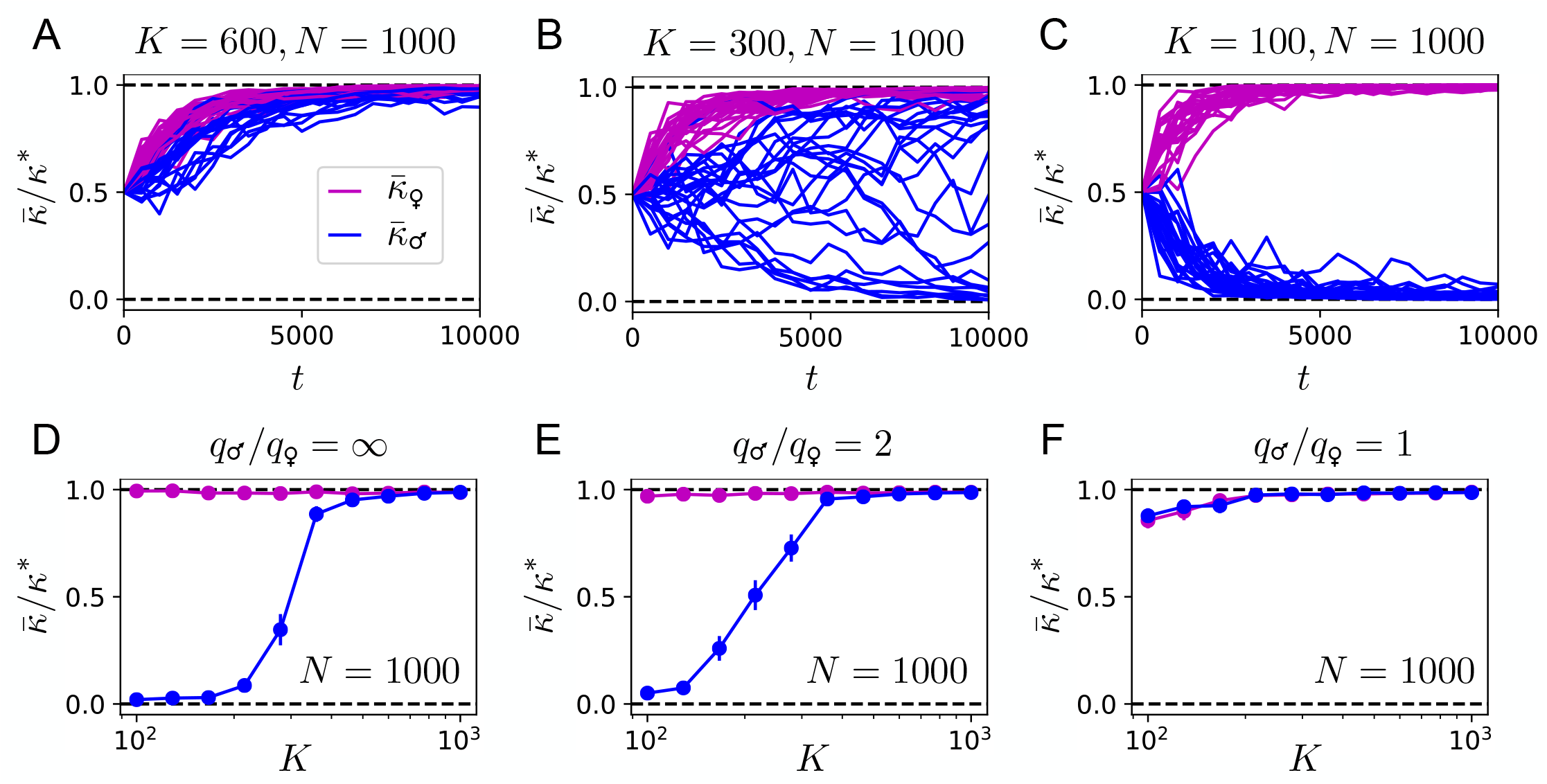
**A-C**. Examples of evolutionary trajectories, showing the mean values of the reaction norms *κ*_♀_ (in pink) and *κ*_♂_ (in blue) for 25 different populations as a function of the number *t* of generations, starting from populations with *κ*_•_ = *κ*^*^/2 for all individuals. In presence of large mating groups (A) monomorphic plasticity evolves where male reaction norms are maximal while in presence of small mating groups (C), dimorphic plasticity evolves where male reaction norms are minimal. In intermediate cases (B), male reaction norms may take arbitrary values. *K* indicates the maximal size of the mating groups and *N* the total population size. **D**. Mean value of the reaction norms *κ*_♀_ and *κ*_♂_ after *T* = 10^4^ generations (averaged over 25 populations) showing how the evolution of dimorphic plasticity depends on the maximal group size *K* of the mating groups (displayed in log-scale). As in panels A-C, the simulations are performed under the hypothesis that fertility is only limited by females (*q*_♂_ = ∞). **E**. If instead of assuming that each male can participate in an arbitrary number of mating events we assume that they can participate in at most twice as many mating events as females (*q*_♂_ = 2*q*_♀_), qualitatively similar results are obtained. **F**. If males and females are subject to the same limitations in fertility (*q*_♂_ = *q*_♀_), however, the symmetry between the two sexes prevents any sexual dimorphism to evolve.

### C. Two-fold cost of males

Our results show that dioecious populations can benefit from sexual dimorphism in plasticity. This benefit is quantified by a supplement of growth rate given by ΔΛ = Λ(*κ*_♀_ = *κ*^*^, *κ*_♂_ = 0) − Λ(*κ*_♀_ = *κ*^*^, *κ*_♂_ = *κ*^*^). Compared to monoecious populations, however, dioecious populations incur a two-fold cost per generation [7], which corresponds to a growth rate difference of ln 2 (Methods). This cost is therefore offset if ΔΛ > ln 2. In general, ΔΛ increases with *c*, the extent of directional selection, and offsetting the two-fold cost of dioecy through dimorphic plasticity therefore requires *c* to exceed a threshold *c*^*^ (Fig. 4A). This is not, however, the only constraint: the survival rate of non-plastic individuals, which decreases with increasing *c*, must also be taken into account. With the optimal reaction norms *κ*_♀_ = *κ*^*^ and *κ*_♂_ = 0, this corresponds to the fraction *χ*_♂_ of males reaching maturation becoming smaller and smaller as *c* becomes larger. Given a population size *N*, it is however necessary to have *χ*_♂_*N* > 1 or no male is expected to be available for mating with females [5]. We find that provided that *κ*^*^ is sufficiently large and 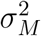 sufficiently low, the survival rate *χ*_♂_ of non-plastic males taken at *c* = *c*^*^ is of the order *χ*_♂_ ∼ 1/2 (Fig. 4B), thus indicating that conditions exist where the two-fold cost of males can be overcome through dimorphic plasticity even in populations of moderate sizes.

**FIG. 4:**
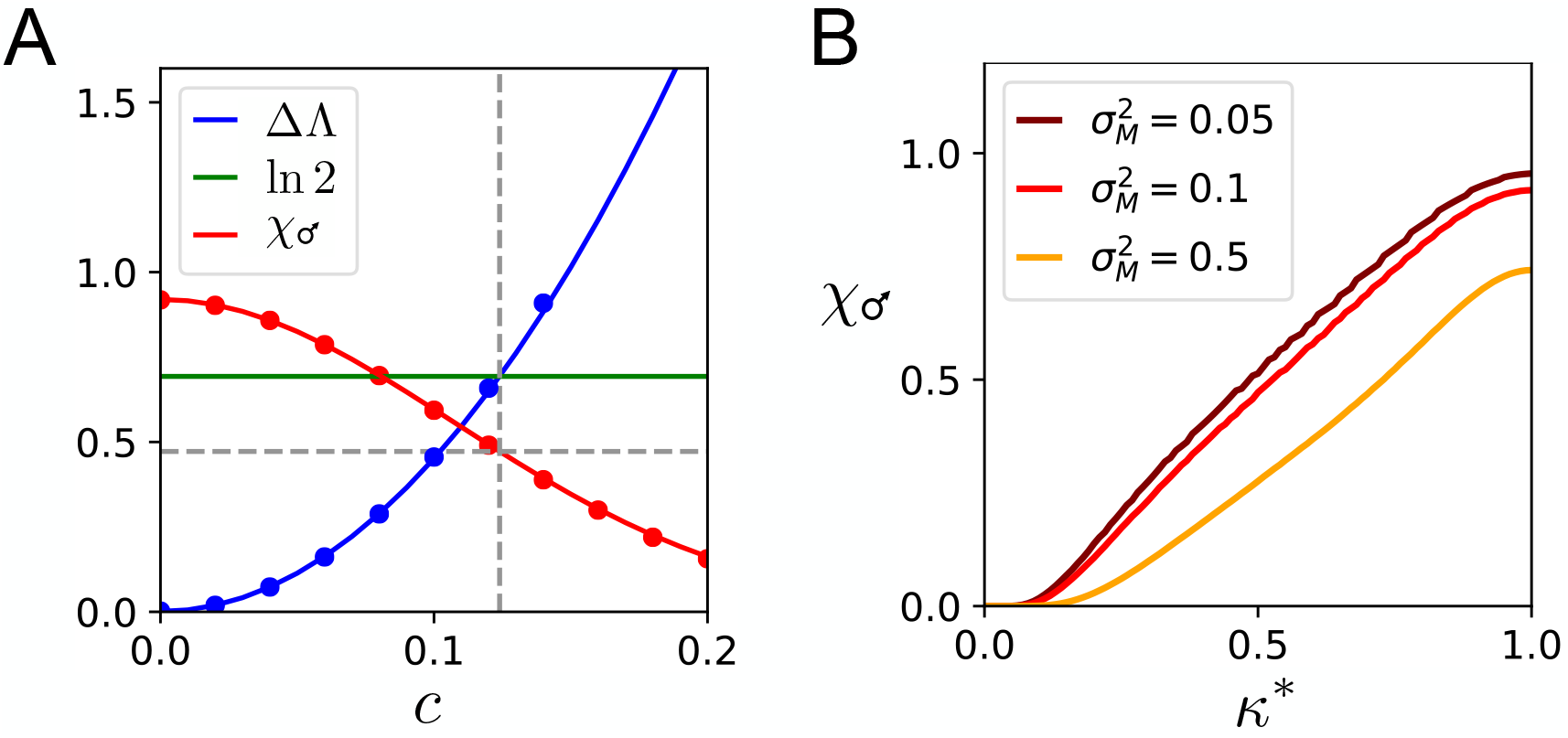
**A**. Gain in growth rate from sexual dimorphism in phenotypic plasticity, ΔΛ = Λ(*κ*_♀_ = *κ*^*^, *κ*_♂_ = 0) − Λ(*κ*_♀_ = *κ*^*^, *κ*_♂_ = *κ*^*^), as a function of *c*, the rate at which the selective pressure is varying (in blue). The vertical dashed line marks the value *c*^*^ above which this difference exceeds the two fold cost of males, ln 2 (in green). The fraction *χ*_♂_ of surviving males at each generation is indicated in red. The lines are analytical calculations and the dots results from numerical simulations. **B**. Fraction *χ*_♂_ of surviving males as a function of the maximal achievable reaction norm *κ*^*^ when considering *c* = *c*^*^, shown here for three different values of the mutational variance 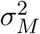. For instance, with 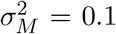 and *κ*^*^ = 0.5 (the values used in all other figures), *χ*_♂_ ≃ 0.5, which indicates that half of the males are expected to reach maturation at any given generation, a survival rate that is easily sustained in finite-size populations.

### D. Greater male variability

As the phenotypic variance (the variance of *ϕ* among mature individuals of a same generation) is a decreasing function of reaction norms, non-plastic males have higher phenotypic variance than plastic females (Methods). We assumed, however, that all members of a same generation experience the same environment *x*_*t*_. A more realistic assumption is that they experience a common average environment *x*_*t*_ with fluctuations that are normally distributed with a finite variance 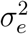 (fluctuations in the mutational bias, on the other hand, simply amount to a larger value of 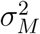). Considering that each individual experiences a particular environment *y*_*t*_ = *x*_*t*_ + *ϵ* with 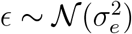 has two implications. First, it changes the probability to reach maturation, which becomes 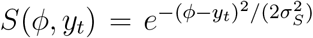. Second, it changes the genotype-to-phenotype map, which becomes *ϕ* = (1 − *κ*_•_)*γ* + *κ*_•_*y*_*t*_ + *ζ* with 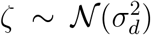, where 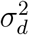 represents the contribution to the developmental variance that is independent of the environment. This more general model can, however, be mapped to the previous model by considering a sex-specific effective developmental variance, 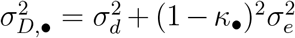. Our results are robust to this generalization, including the fact that dimorphic plasticity can evolve where males have a more limited phenotypic plasticity associated with a greater phenotypic variability (Fig. S2).

## IV. DISCUSSION

We presented a mathematical model for the evolution of phenotypic plasticity in dioecious populations which led us to the following conclusions:

1. Sexual dimorphism in phenotypic plasticity can be optimal for long-term population growth when selection is directional, either because of a directionally varying environment or a systematic mutational bias. The root of the asymmetry is the limited fertility of one of the two sexes, conventionally females. The optimum then corresponds to females being maximally plastic (with a high reaction norm) and to males being non-plastic (with no reaction norm).
2. The evolution of plasticity in panmictic populations leads to maximal plasticity in the two sexes, i.e., a suboptimal long-term growth rate, which is interpreted as arising from sexual selection or, more precisely, from selection on male viability as our model assumes no female choice.
3. Limiting mating to within groups of sufficiently small sizes can counter-act sexual selection and enable the evolution of sexual dimorphism in phenotypic plasticity.
4. Sexual dimorphism in phenotypic plasticity can confer an evolutionary benefit of dioecy over monoecy despite the two-fold cost of males.
5. As a smaller reaction norm implies larger phenotypic variance, and even more so when environmental differences are present within a generation, the sexual dimorphism in phenotypic plasticity that evolves implies a greater male variability.

The first point was demonstrated in previous works [4, 5] but the conditions under which dimorphic plasticity may evolve was not previously established. Here we show that limiting mating to within groups is essential to counter-act the adverse effect of intra-male competition and thus resolve a conflict between levels of selection in favor of the highest, population level. At variance with most other explanations of sexual dimorphism, sexual dimorphism in phenotypic plasticity thus arises in our model not because but in spite of sexual selection.

A greater male variability is commonly observed in many species, as for instance reviewed in [15]. Several alternative explanations have been proposed [16], including scenarios involving a stronger selection on males [17], consistent with our results where plasticity evolves to buffer selection on females. Other scenarios have also been proposed to explain how two sexes may provide an adaptive advantage that offsets the two-fold cost of males [19–25], which all elaborate on the informal idea that the two-fold cost of males is offset if reproducing males are twice as fit as reproducing females [26]. All these scenarios rely on the same fundamental asymmetry, differential variance in mating success. As most previous models on the subject [20–22], we have only shown here that sexually dimorphic plasticity may provide an adaptive advantage for dioecy over monoecy but not examined the conditions under which dioecy may evolve from monoecy. This would require analyzing the conditions under which a dioecious mutant can invade a monoecious population. While this question is beyond the scope of the present work, we note that the few available models demonstrating the evolution of dioecy either do not consider a twofold cost of males, e.g. [27], or assume an additional asymmetry between the two sexes [23, 24], namely an asymmetry in the strength at which the two sexes are selected, with males being subject to a stronger selection than females. In our model, the only assumed asymmetry is in differential reproductive investment – the defining distinction between males and females. We find that the two sexes evolve to experience differential selective pressure, but this is an emergent feature arising from the evolution of sexual dimorphism in phenotypic plasticity, not an a priori assumption. Our scenario thus resembles the formal model of Holmes et al [25] where males similarly pay most of the cost of adaptation and it even more closely resembles the informal theory of Geodakyan [18], although none of these previous studies anticipated the antagonistic effect that sexual selection may have. This effect imposes a strong constraint: mating must occur within groups of sufficiently closely related individuals for dimorphic plasticity to evolve as a plastic male within a group of non-plastic males has always a higher chance of reproducing. More generally, we note that the same mechanism by which sexual dimorphism in phenotypic plasticity evolves through group/kin selection in our model is at play in other models for the evolution of biological symmetry breaking [28]. This bolsters the hypothesis that the evolution of two sexes might instantiate a more general evolutionary phenomenon [29].

## Acknowledgements

We thank Nobuto Takeuchi for stimulating discussions.

## Methods

### Dynamics for large unstructured populations

In the limit of large population sizes and in absence of group structure, the long-term growth rate Λ of the population can be obtained analytically by solving a recursion for the density *n*_*t*_(Γ) of newborn individuals at generation *t* with genotype Γ = (*γ, y*_♀_, *y*_♂_), where 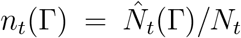 if 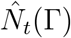 is the mean number of individuals born at generation *t* with genotype Γ and 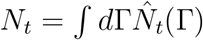 the mean total population size.

First, the mean number 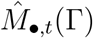 of individuals of sex • = **♀** or **♂** that reach maturation is

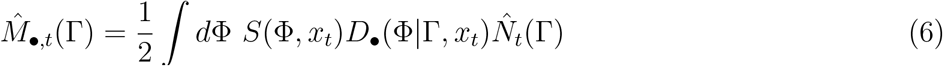

where *D*_•_(Φ|Γ, *x*_*t*_) is the probability that an individual of sex • and genotype Γ acquires a phenotype Φ = (*ϕ*, •, *κ*) and *S*(Φ, *x*_*t*_) is the probability that it survives to reach maturation, given an environment *x*_*t*_. The factor 1/2 accounts for the fact that sexes are assigned randomly with same probability. This may be rewritten in terms of densities as

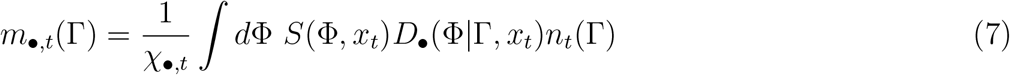

where 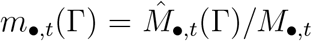 with 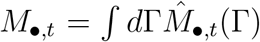 is the density of mature individuals of sex • at generation *t*, and where *χ*_•,*t*_ represents the fraction of individuals of sex • reaching maturation, which is given by

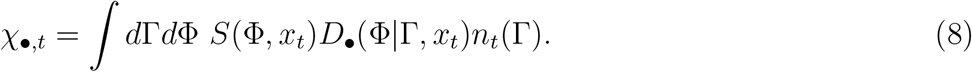

Second, random mating between individuals of the opposite sex lead to a density of newborn individuals with genotype Γ at generation *t* + 1 given by

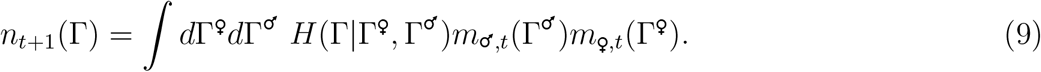

where *H*(Γ|Γ^♀^, Γ^♂^) is the probability that a couple with genotypes Γ^♀^ and Γ^♂^ produces an individual of genotype Γ.

The total number of mating events at generation *t* is min(*q*_♀_*M*_♀,*t*_, *q*_♂_*M*_♂,*t*_) where 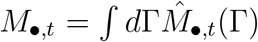 is the mean total of individuals of sex • that reach maturation. Growth is thus limited by females if *q*_♀_*χ*_♀,*t*_ < *q*_♂_*χ*_♂,*t*_ and by males otherwise. If for instance females are the limiting sex, the growth increase at generation *t* is

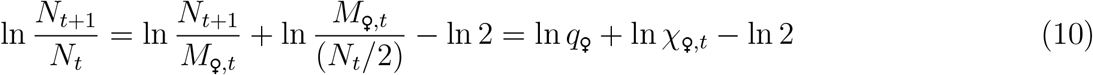

and the same expression with **♂** instead of **♀** if males are the limiting sex.

Finally, for directionally varying selective pressures, *χ*_♀,*t*_ and *χ*_♂,*t*_ asymptotically reach constant values *χ*_♀_ and *χ*_♂_ so that the long-term population growth is given by

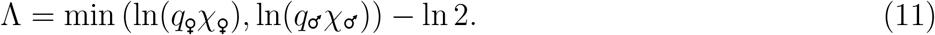

For monoecious (hermaphroditic) populations, the same formulas apply but with *m*_♀,*t*_(Γ) = *m*_♂,*t*_(Γ) = *m*_*t*_(Γ) and without the term ln 2 in Eq. (11). This term corresponds to the two-fold cost of males when fertility is limited by females and, more generally, to the two-fold cost of dioecy.

In our model, all the kernels are Gaussian, that is, if we denote Gaussian functions by 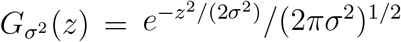,

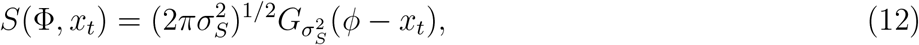

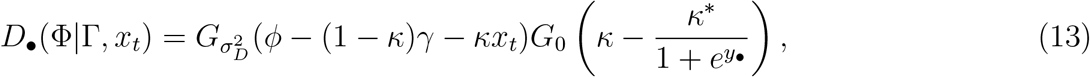

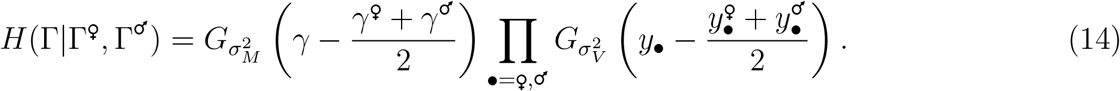

Here *G*_0_(*z*) = *δ*(*z*) represents Dirac delta function.

### Analytical calculations

We consider here the model with 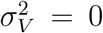 and fixed values of *κ*_•_ for which the genotypes Γ And phenotypes Φ reduce to scalar variables *γ* and *ϕ*. We treat a more general case where *ϕ* = (1 − *κ*_•_)*γ* + _*D*_ *κ*_•_*x*_*t*_ + *ζ* with 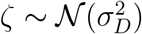 where λ_•_ may differ from 1 − *κ*_•_ so that

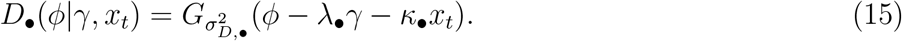

For the sake of generality, we also allow for the possibility that 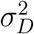, *c*_*M*_ and 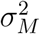 are sex-dependent. We otherwise adhere to the definition of the model given in the main text. The derivation below follows the solution that we previously provided for a more general model [5].

As the densities *n*_*t*_(*γ*) and *m*_•,*t*_(*γ*) are asymptotically normally distributed we make the following Gaussian Ansätze to solve Eqs. (7) and (9):

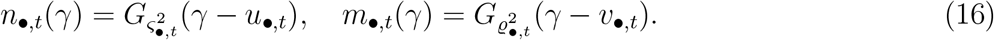

This leads to

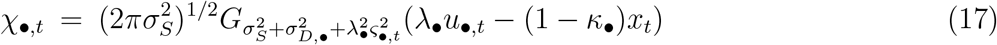

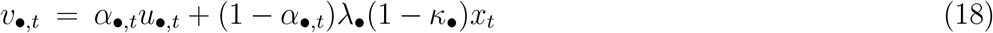

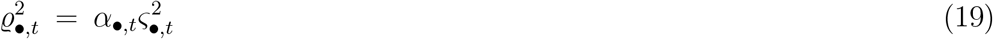

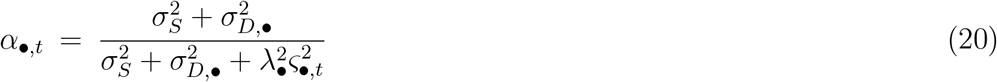

and

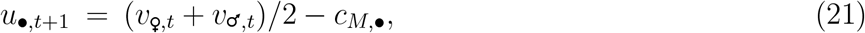

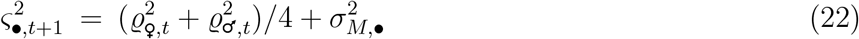

To accommodate for sex-specific genetic parameters, it is convenient to introduce

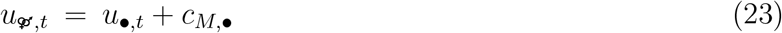

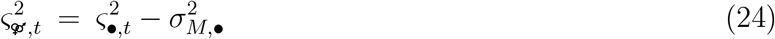

which are independent of • and follow the recursion

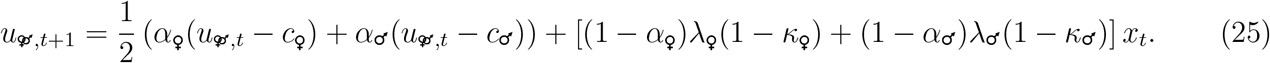

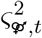 reaches a fixed point 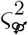 when *t* → ∞ that is solution of the cubic equation

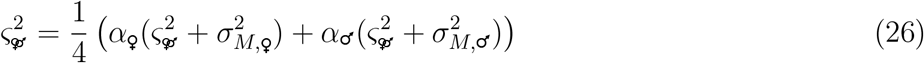

with

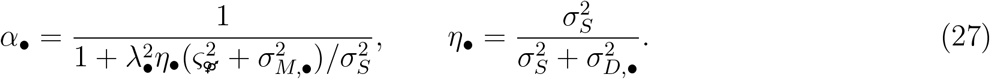

Finally, if we assume for instance that females are the limiting sex we have Λ = ln(*q*_♀_*χ*_♀_/2) with

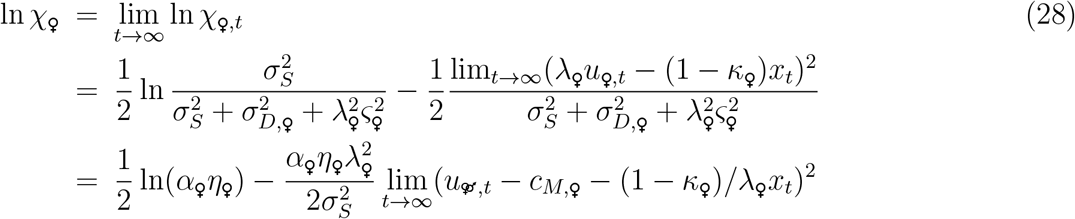

Here note that we can always assume *c*_*E*_ = 0 by redefining *c*_*M*,9_ as *c*_*M*,9_ + (1 − *κ*_9_)/*λ*_9_*c*_*E*_. We are therefore left with the evaluation of 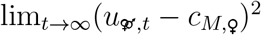 where 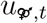 is given by

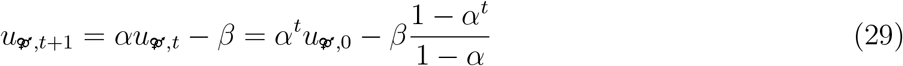

with

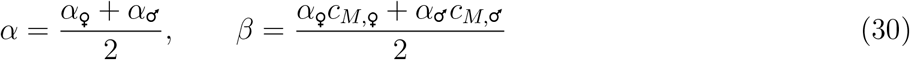

so that

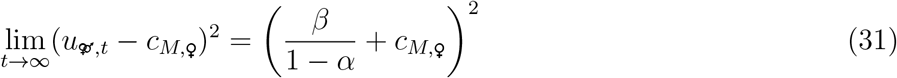

Finally, considering *c*_*M*,♀_ = *c*_*M*, ♂_ = *c*_*M*_ and a non-zero *c*_*E*_, we obtain Λ as given by Eq. (11) with

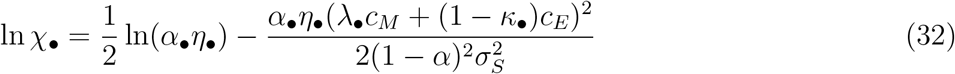

### Phenotypic variance

If *n*_*t*_(*γ*) is normally distributed, the phenotypes *ϕ* of mature individuals of sex • are also normally distributed with a distribution *m*_•,*t*_(*ϕ*) given up to a normalizing factor by

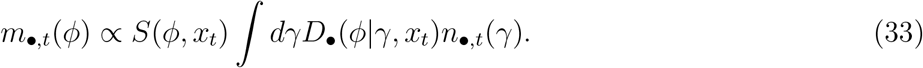

The phenotypic variance is therefore

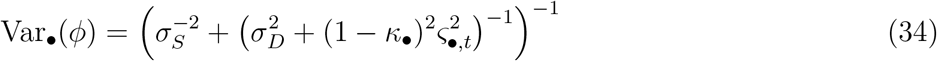

When the mutational variance 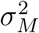 is independent of the sex, the genetic variances 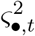 reach a fixed point *ς*^2^ that is also independent of the sex •. In this case, the smaller the reaction norm, the larger the phenotypic variance. In presence of intra-generation environmental fluctuations, 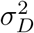 is replaced by 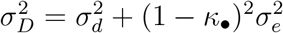, which does not change this conclusion.

### Numerical simulations

As in our previous works [5], numerical simulations are performed with a population of fixed size *N*. Genotypes Γ are initially identical for all members of the population and a phenotype Φ is assigned to each individual based on its genotype Γ and the environment *x*_*t*_ as described in the main text. Each individual is then selected with probability 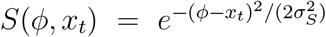. At generation *t*, this results in *M*_♀,*t*_ mature females and *M*_♂,*t*_ mature males. The total number of mating events is then *N*_*t*_ = min(*q*_♀_*M*_♀,*t*_, *q*_♂_*M*_♂,*t*_). If for instance *q*_♀_*M*_♀,*t*_ < *q*_♂_*M*_♂,*t*_, each female is involved in exactly *q*_♀_ mating events and males are sampled at random without replacement in the set of size *q*_♂_*M*_♂,*t*_ where each male is represented *q*_♂_ times or, if *q*_♂_ = ∞, sampled at random with replacement in the set of males. The same procedure is applied with the two sexes playing opposite roles if *q*_♀_*M*_♀,*t*_ *> q*_♂_*M*_♂,*t*_. Each mating event leads to a newborn that inherits a genotype Γ based on the genotypes Γ^♀^ and Γ^♂^ of its parents following the rules given in the main text. The population at the next generation *t* + 1 is then obtained by randomly sampling with replacement *N* of the *N*_*t*_ newborns. *N* may be larger than *N*_*t*_, but we restrict the study to parameters where *N*_*t*_ > 0 at each generation *t*. After *T* generations, the long-term growth rate is estimated as 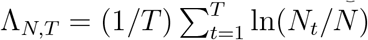.

### Extension to soft selection and migration

Regulation to conserve a constant population size is performed by resampling *N* individuals irrespectively of their group, which corresponds to a regulation at the global population level known as “hard selection” [30, 31]. An opposite limit is when density regulation occurs at the level of each group independently, also known as “soft selection”. This can be implemented in our model by changing the way in which each of the *N* individuals is sampled, namely by first drawing a group uniformly at random before drawing an individual at random in this group. More generally, we can interpolate between these two extreme cases by introducing a parameter 0 *< u <* 1 where *u* = 0 describes hard selection only and *u* = 1 soft selection only. To this end, each of the *N* individuals forming a new generation is obtained as follows: with probability 1 − *u*, a random individual is drawn irrespectively of its group while with probability *u*, a random group is first drawn, from which a random individual is then drawn. Results obtained by varying *u* are shown in Fig. S3A.

We can also introduce exchange between mating groups by having a non-zero probability *m* that each individual migrates to a new group. The results obtained by varying *m* when making this exchange after density regulation and when choosing the new group in proportion of the size are shown in Fig. S3B. Variants of this procedure are possible which give qualitatively similar results. Finally, soft-selection (*u* > 0) and migration (*m* > 0) can both be present as in Fig. S3C.

## Codes and data availability

All codes and data used to generate the figures are available at https://github.com/orivoire/evosexdimo.

---

**FIG. S1:**
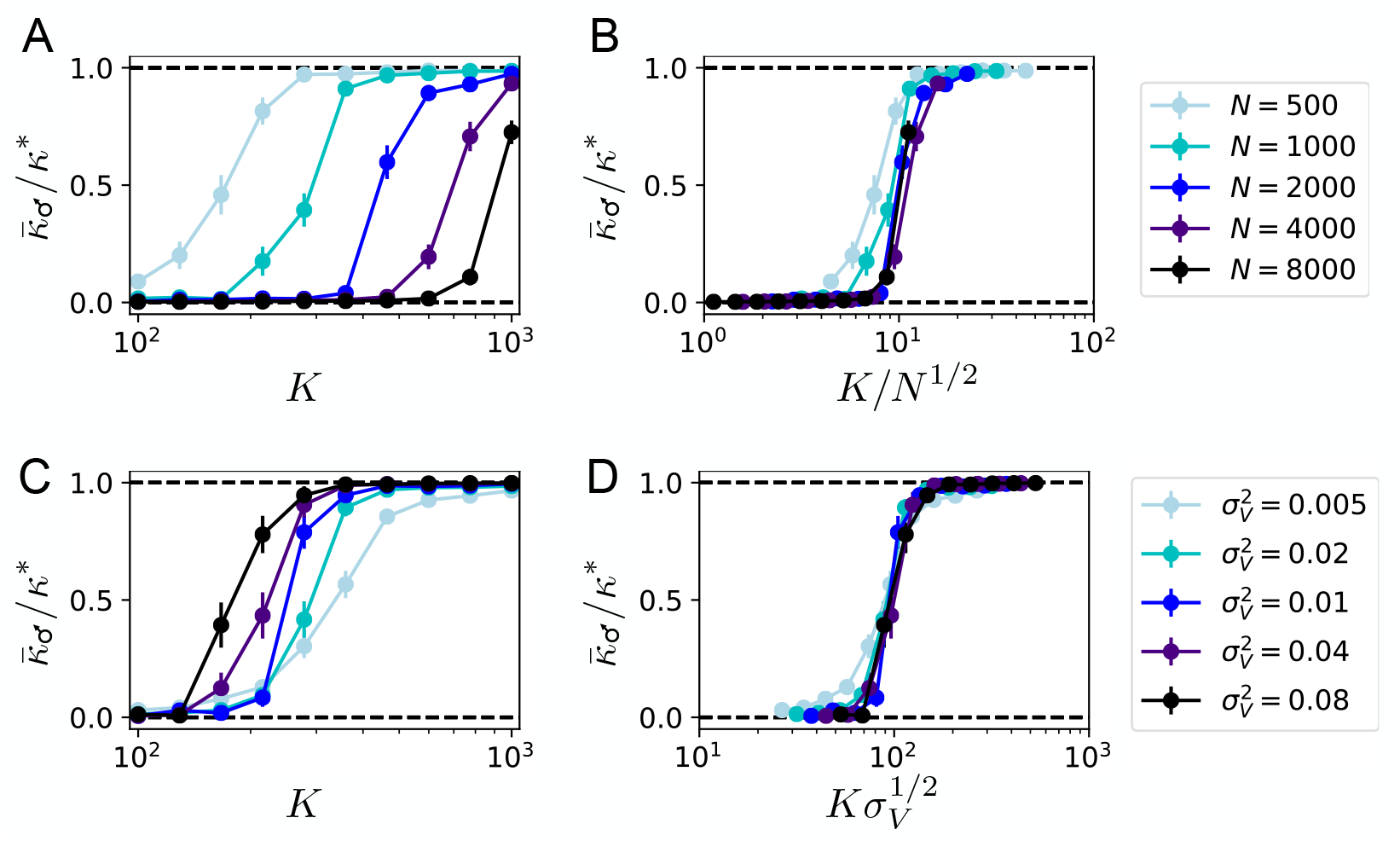
Scaling of the maximal group size below which dimorphic plasticity evolves – The calculations performed to obtain Figure 3D are repeated here with either different total population sizes *N* (A-B) or different mutational variances for the modifier genes 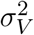 (C-D). **A**. As the total population size *N* increases, the maximal group size *K*^*^ below which dimorphic plasticity evolves increases. **B**. This maximal group size *K*^*^ scales as *K*^*^ ∼ *N* ^1/2^ for large values of *N*. **C**. As the mutational variance 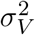 increases, the maximal group size *K*^*^ below which dimorphic plasticity evolves decreases. **B**. This maximal group size *K*^*^ scales as 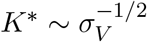. Only average male reactions norms are shown here as female reaction norms evolve to maximal values in all cases, and results are averaged over 25 independent populations as in Figure 3D.

**FIG. S2:**
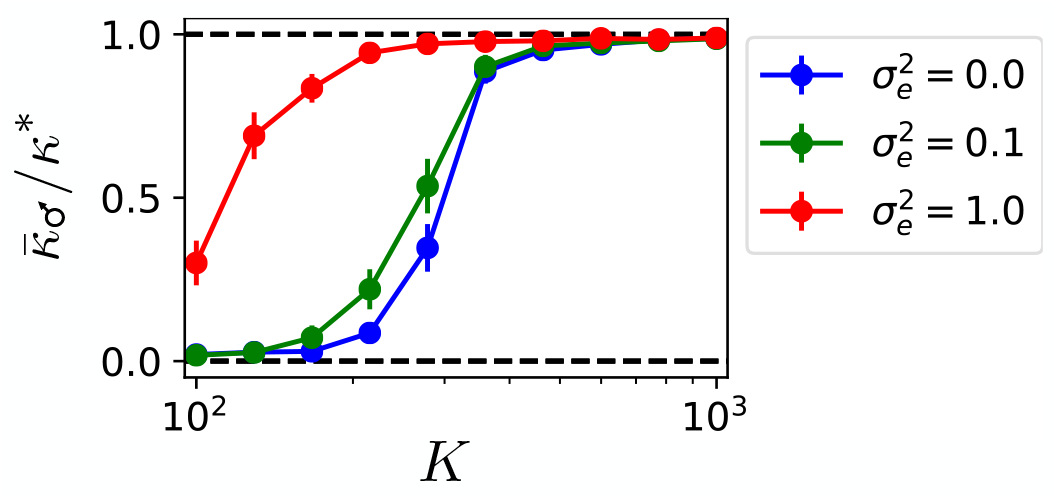
Evolution of plasticity in presence of intra-generation environmental fluctuations – The parameters and calculations are as in Figure 3D, except that each individual at generation *t* experiences a local environment *y*_*t*_ = *x*_*t*_ +*t:* where *t:* is normally distributed with zero mean and variance 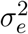. As a consequence, the probability to reach maturation is *S*(*ϕ, y*_*t*_) instead of *S*(*ϕ, x*_*t*_) and the phenotypes are related to genotypes by *ϕ* = (1 − *κ*_•_)*γ* + *κ*_•_*y*_*t*_ instead of *ϕ* = (1−*κ*_•_)*γ* +*κ*_•_*x*_*t*_. Here we consider *c*_*M*_ = 0 and therefore *c* = *c*_*E*_. The results for 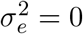 correspond to those of Figure 3D. For sufficiently small 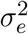, the results are qualitatively unchanged: sexual dimorphism in plasticity evolves when mating occurs within groups of sufficiently small size *K*. Large values of 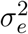, however, tend to disfavor the evolution of dimorphic plasticity.

**FIG. S3:**
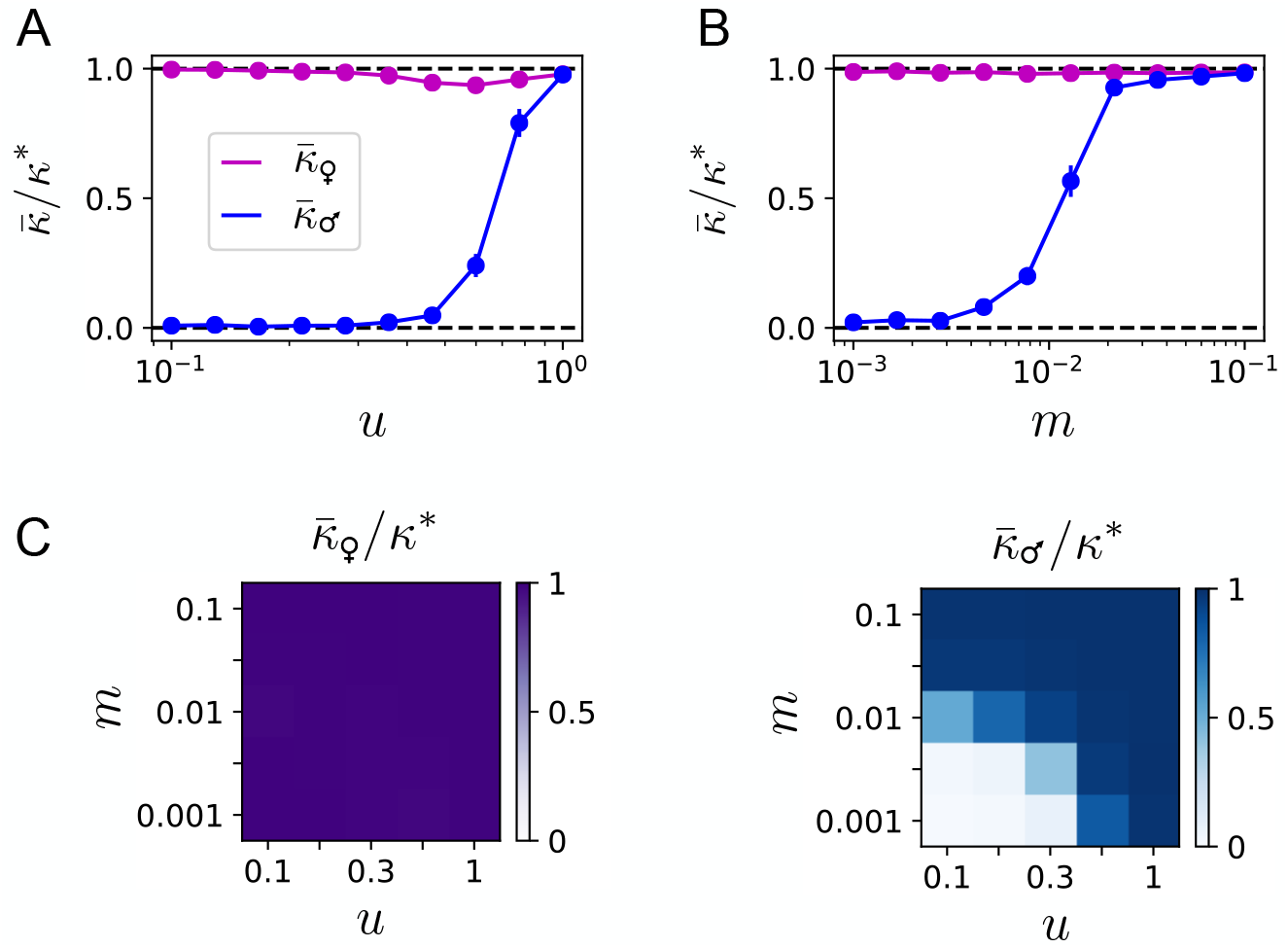
Extension of the model to partial soft selection and to migration between mating groups – The model analyzed in Figure 3C, with *K* = 100 and *M* = 1000, is generalized to include soft-selection (*u* > 0) or/and migration (*m* > 0) (Methods). **A**. The parameter *u* is introduced to interpolate between a regulation of density exclusively at the population level (hard selection, *u* = 0 as in the main text) and a regulation of density exclusively at the group level (soft selection, *u* = 1). Sexual dimorphism in phenotypic plasticity evolves provided *u* is small enough, i.e., provided sufficient density regulation at the global level takes place. **B**. The parameter *m* is introduced to interpolate between strictly independent groups (no migration, *m* = 0 as in the main text) and a situation where group identities are completely randomized at each generation (probability *m* = 1 of migrating). Sexual dimorphism in phenotypic plasticity evolves provided *m* is small enough, i.e., provided exchanges between groups are small enough. **C**. More generally, sexual dimorphism in phenotypic plasticity with high female plasticity 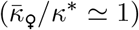 and small male plasticity 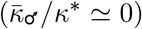 evolves when the two parameters *u* and *m* are both sufficiently small. Note that *u* and *m* are represented on a logarithm scales.

